# Evolve and resequence provides a granular view of micro-evolution under different mating systems in *Mimulus guttatus*

**DOI:** 10.64898/2025.12.10.693597

**Authors:** Sharifu K. Tusuubira, Luis Javier Madrigal-Roca, Keely Brown, John K. Kelly

## Abstract

We performed 10 generations of experimental evolution in *Mimulus guttatus* and measured genome-wide change in replicated populations that were compelled to reproduce entirely by self-fertilization, entirely by outcrossing, or by a mixture of the two. We developed a novel testing framework based on ancestral haplotype inference to locate mating system loci. Our results confirm several outstanding theoretical predictions: Selfing populations showed increased homozygosity, widespread hitch-hiking, and higher stochastic changes in allele frequencies compared to outcrossing populations. Despite this variability, approximately 20 genomic regions (QTLs) demonstrated parallel evolution across treatments. We identified candidate genes within QTLs using RNA sequencing data from the ancestral lines. In several instances, we found closely linked candidate genes, suggesting that by inhibiting recombination inbreeding can allow for selection on favorable gene combinations. We observed a general down-regulation of candidate genes in selfing populations, mirroring known transcriptome differences between established selfing and outcrossing sister species. This suggests that gene expression is a significant component of the “selfing syndrome.”

## Introduction

The evolution from outcrossing to predominant self-fertilization (selfing) is one of the most frequent transitions in the history of flowering plants and has been a focus of sustained investigation since Darwin (1876). Selfing species often differ strikingly from closely related outcrossing taxa in both morphology (Ornduff 1969) and genetic composition (Wright *et al*. 2008; Zhang *et al*. 2022). It is an open question as to how much of this divergence is driven by natural selection in the initial transition to selfing (based on standing variation within the ancestral outcrossing species) versus the fixation of *de novo* mutations after selfing has been established. This paper considers the initial stage of this process focusing on two key aspects. First, how rapidly does selfing transform the genomic landscape through increased homozygosity? Given increased stochasticity and a reduced efficacy of recombination, can selection effectively recruit alleles from across the genome into multi-locus genotypes that are both viable and reproduce efficiently by selfing? Second, can we map genes that respond to selection driven by mating system change? Mapping enables a comparison of genetic features between standing variation and the loci that distinguish divergent taxa.

There is increasing evidence that outcrossing populations *can* rapidly respond to selection for increased selfing in response to a partial or complete loss of pollinator service. Temporary declines in pollinators have probably always been an important selective agent on flowering plants (Brys and Jacquemyn 2011), but human-driven changes make this a major focus of current research (Potts *et al*. 2016). Resurrection experiments in *Ipomoea purpurea* and *Viola arvensis* have demonstrated contemporary phenotypic evolution of mating system traits in response to pollinator declines (Bishop *et al*. 2023; Acoca-Pidolle *et al*. 2024). Previous work on *Mimulus guttatus* showed that experimental populations maintained without pollinators rapidly diverged from initially equivalent replicate populations maintained with pollinators (bumblebees), both in terms of morphology and fitness components (Bodbyl Roels and Kelly 2011), as well as genome-wide sequence variation (Busch *et al*. 2022). We here describe an experimental evolution study of the same species, but using an experimental design optimized to provide a more detailed picture of genome-wide changes and the loci under selection.

Geneticists have successfully mapped loci with “Mendelian” effects on the mating system. For instance, transitions from bee to hummingbird pollination in *Penstemon* spp. are associated with a shift from blue to red flowers. These changes are caused by the functional inactivation of a specific anthocyanin pathway gene; loss-of-function mutations to this gene have fixed independently in many hummingbird pollinated lineages (Wessinger and Rausher 2014). Crosses between genetically self-compatible and self-incompatible species routinely produce mendelian ratios for compatible/incompatible plants in F_2_ panels, e.g. (Slotte *et al*. 2012; Koseva *et al*. 2017; CarrÉ *et al*. 2021) and these QTLs have been resolved to the causal gene (or genes) in species where the S-locus has been characterized (Slotte *et al*. 2012; Li and Chetelat 2015), but also see (Li *et al*. 2023). With a few notable exceptions (Chen *et al*. 2007; Sicard *et al*. 2016), there has been less progress on the fine-scale genetic characterization of quantitative trait changes altering mating system. Most mapping studies have examined crosses between divergent species, e.g. *Mimulus* (Lin and Ritland 1997; Fishman *et al*. 2002), *Capsella* (Sicard and Lenhard 2011; Slotte *et al*. 2012), and *Solanum* (Bernacchi and Tanksley 1997). Inter-species crosses map changes that both occurred during transition to selfing and mutations that fixed after that transition. Sicard *et al*. (2016) resolved a mating system QTL to the causal gene and showed that an interspecific difference (the allele fixed in the selfing species) was recruited from standing variation in the ancestral outcrossing population.

Our experimental design is illustrated by Fig 1. We started by intercrossing 187 fully homozygous and genome sequenced lines derived from a single natural population of *Mimulus guttatus*. This population (Iron Mountain, hereafter IM) reproduces primary by outcrossing, on average about 90% (Monnahan *et al*. 2021). Seeds from the Ancestral population were randomly sorted into 20 experimental populations, and over the next 10 generations, these populations independently evolved within three different treatments: all reproduction by outcrossing, all by self-fertilization, or a by a mixture of selfing (90%) and outcrossing (10%). There was substantial phenotypic evolution over the ten generations with divergence between mating system treatments in rate of development (days to flower), flower morphology (corolla dimensions and stigma-anther separation), and reproductive allocation, particularly the ability to set seed without pollinators (Tusuubira and Kelly 2024). For the present study, we genome sequenced pooled samples from each descendant population to identify loci underpinning these changes, i.e. loci that exhibit consistent differentiation between mating system treatments. This design exploits the idea that we can accurately track the frequencies for entire haplotypes within evolving populations. This enables accurate estimation of allele and haplotype frequencies, tracking of haplotype dynamics at the whole chromosome level, testing for a multi-allelic response to selection, and determining whether genomic regions under selection exhibit distinct patterns of gene expression.

**Figure 1.**
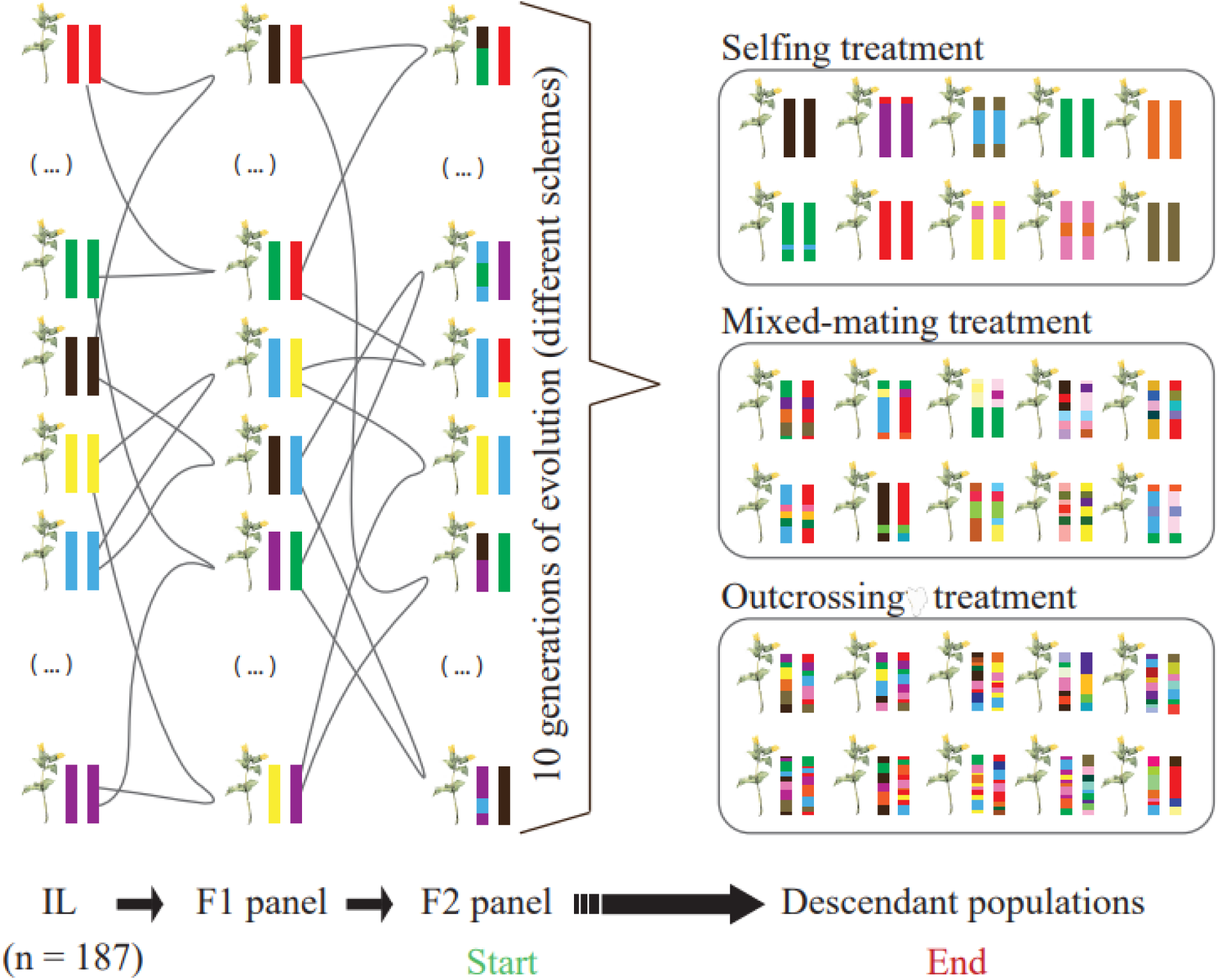
A diagram of this experiment with different colors coding the 187 ancestral lines that were intercrossed to create the Ancestral population. Chromosomes in Outcrossing descendant populations are predicted to be more finely intermixed in ancestry than Selfing population descendants.

High sequencing depth is usually required to accurately estimate allele frequencies in pooled samples, but this is not true if populations are derived from ancestral haplotypes of known sequence (Kessner *et al*. 2013; Tilk *et al*. 2019). The plants in an evolved population will be a mosaic of ancestral haplotypes (Fig 1), but within any localized region, a plant is likely to carry the DNA from only one or two ancestral lines. Therefore, we can leverage information across neighboring SNPs to infer the contribution of each ancestral haplotype to the current population. These contributions change from window to window along a chromosome owing to recombination. Since the genotype of all ancestral lines are known, the inferred ancestral line frequencies can be accurately estimated at any SNP scored in the ancestral haplotypes (Tilk *et al*. 2019; Wu *et al*. 2025). One purpose of this paper is to develop and apply multi-allelic testing methods based on the estimated ancestral haplotype frequencies and test how these frequencies differ among mating system treatments. Founding the experiment from fully sequenced lines also enables the discovery of multi-allelic variation. For practical reasons, evolve and resequence experiments have focused on evolution of biallelic SNPs. However, scored SNPs may not be the primary cause of fitness effects. The two alleles at a SNP may often exhibit imperfect associations (linkage disequilibria) with unscored polymorphisms that are the actual targets of selection. In this circumstance, haplotype dynamics may be more informative than SNP dynamics (Barghi and SchlÖtterer 2019; Michalak *et al*. 2019), which is particularly true when more than two functionally distinct alleles segregate in a population. This appears to be very common for QTLs in both plants and animals (Buckler *et al*. 2009; King *et al*. 2014). Within the IM population of *M. guttatus*, most of the genetic variation in gene expression is generated by cis eQTLs that vary as an allelic series with three or more functionally distinct variants (Veltsos and Kelly 2024). With only two alleles at a locus, we might expect natural selection to favor the “selfing-favorable” allele strongly in the Selfing treatment, and more moderately in the Mixed-mating treatment (Fig 1). However, if a mating system locus segregates as an allelic series in the Ancestral population, different alleles could be favored in each treatment.

Another advantage of founding from sequenced lines is that if we measure evolutionary change in terms of the frequencies of ancestral haplotypes, selection responses to can be directly related to any other measured feature of these haplotypes. Inbred lines are “immortal genotypes” replicable across different experiments. The IM lines have previously been characterized for gene expression variation in flower buds (Brown and Kelly 2021). Thus, we can directly relate evolutionary changes in the present experiment to genetically based differences in gene expression demonstrated previously. Consider a locus under selection such that a particular set of haplotypes consistently increase within the Selfing but not Outcrossing populations. We can ask if those same ancestral lines exhibit elevated or reduced gene expression driven by variants within the indicated genomic region. Importantly, there is no need to force these comparisons into binary categories. Within each locus/window, each ancestral line yields continuous values for the change in frequency within each replicated population of each mating system treatment and for the relative levels of expression of genes in that window. We can effectively integrate this information, testing for associations, by treating the ancestral haplotype as the unit of replication. Finally, by “tagging” ancestral haplotypes, this experiment provides a granular, chromosome level view of microevolution following pollinator loss (Fig 1). We predict sustained genetic interchange among individuals in the Outcrossing treatment, and after 10 generations, descendant genomes will be highly refined combinations of many ancestral genomes. As a consequence, favorable alleles should be distributed across many genetic backgrounds and hitch-hiking effects closely localized to fitness determining loci (Maynard Smith and Haigh 1974). In contrast, homozygosity should increase rapidly in the Selfing populations with a corresponding reduction in the efficacy of recombination. We predict that the joint action of selection and selfing will reduce genetic variation far beyond the predicted 50% reduction in effective population size (Pollak 1987). With complete selfing, the population is a collection of competing lineages and hitch-hiking should extend over large chromosomal regions. The primary products of this research are (a) a new statistical testing framework to evaluate multi-allelic responses to selection, (b) a demonstration that adaptation to loss of pollinator service involves recruitment alleles that were rare or at least uncommon in the ancestral population, (c) the identification of candidate genes within selected loci from the combination allele frequency change and previous gene expression / functional experiments, and (d) a chromosome-level view of how mating system and selection can shape the genealogical structure of populations.

## Methods

### Study species and experimental evolution

***—****Mimulus guttatus* (syn *Erythranthe guttata*; 2n = 28; Phrymaceae) is a model plant for genetics and mating systems evolution (Wu et al., 2008). We founded this experiment from genotypes sampled from one natural population located on Iron Mountain in Oregon (44°24′03″N, 122°08′57″W). This population is predominantly outcrossing with little internal structure (Willis 1993; Sweigart *et al*. 1999). Starting with seed from thousands of unrelated field plants, we initiated inbred line formation by successive rounds of self-fertilization, a.k.a. single seed descent (Willis 1999b; Kelly 2003). Among several hundred lineages that survived at least 8 generations of selfing (the great majority went extinct due to inbreeding depression), we obtained whole genome sequences for 187 highly homozygous lines (Troth et al., 2018). For this experiment, we grew each line to maturity and randomly paired and crossed them to make F_1_ families. We then randomly intercrossed F_1_ plants to produce F_2_ seeds and combined F_2_ seed from all families into a common pool (our “Ancestral population”). We sampled 30mg of seed from the Ancestral population to establish each of 20 experimental populations: 4 replicates that reproduce only by outcrossing, 8 that reproduce only by selfing, and 8 that reproduced by a mixture of outcrossing (10%) and selfing (90%).

Tusuubira and Kelly (2024) provide a detailed description of the selection experiment and the phenotypic response to selection. Briefly, we started each generation by distributing seed over a 30.5cm x 38.1cm growth arena. Flats were maintained in a pollinator-free environment where all reproduction requires autogamous selfing or hand pollinations. In the Outcrossing treatment, all flowering plants were hand-pollinated using a random sire from the same population on days 24-25 post-seeding. Only seed from these hand-pollinations was used to make the next generation. In the Mixed mating populations, 40 flowering plants were randomly selected to receive hand pollination from another flowering plant on days 26-27. One flower was selected as either a pollen donor or recipient per plant. At the final harvest, we collected the hand pollinated fruits into one envelope and all others (those produced by selfing) into another. To make the founding seeds for the next generation, we combined 3mg from the hand-pollination collection and 27mg from the selfed collection (which enforces the 90% selfing condition for this treatment). In the pure Selfing treatment populations, plants were left undisturbed until the end of the generation and all fruits then collected. While thousands of seeds were in each 30mg propagule, adult population sizes were typically about 400 plants per population owing to natural thinning. Flowering typically began around days 21-23, and after a progressive drought regime, seeds were collected from all plants on days 45-50. This regime mimics field conditions where nearly all plants eventually die of desiccation.

### Sequencing and variant calling within pooled population samples

—After 10 generations of selection, we grew 200 plants from each population for whole-genome sequencing. Each plant contributed equal leaf material (∼10mg) to a combined DNA sample. We extracted DNA from the leaf pool of each population using the cetyltrimethylammonium bromide (CTAB) extraction method (Moreira & Oliveira, 2011; Wang et al., 2012). For the Ancestral population, we made two replicate pools, A1 and A2, each from 200 distinct plants. For each of the 22 DNA pools, the KU genome sequencing core made barcoded libraries using the Illumina DNA Prep kit. We sequenced these libraries using Illumina NovaSeq (S1 option) at the University of Kansas Medical center obtaining slightly more than one billion 100bp single-end reads across all populations.

We mapped sequence reads to the *M. guttatus* IM62v3 reference genome (https://phytozome-next.jgi.doe.gov/) and trimmed the raw reads to remove low-quality bases using FASTP (Chen *et al*. 2018). Reads were aligned using the default settings in BWA (Li and Durbin 2009) yielding a sequence alignment ( “bam”) file for each population. For the ancestral line genomes, we remapped the original sequence data (Troth *et al*. 2018) to the IM62v3 reference genome using BWA, and then obtained a variant call file (“vcf”) for the ancestral lines using mpileup piped to the call command in BCFtools version 1.9 (Danecek *et al*. 2021). We wrote python programs to implement mapping and variant calling. These programs, as well as all those described below, are archived in https://github.com/jkkelly/Mating_system_evolve_resequence. Ancestral haplotype frequencies were obtained using HAFpipe (Kessner *et al*. 2013; Tilk *et al*. 2019; Madrigal-Roca 2025), which takes the bam (populations) and vcf (ancestral lines) files as inputs. For each of the 22 population samples, we obtained ancestral haplotype proportions (187 numbers that sum to one) within 50kb windows. Consecutive windows overlap with a 5kb stagger (0-50kb, 5-55kb, 10-60kb, etc) yielding a total of 61,886 windows over the 14 main chromosomes of *M. guttatus*. These inferred ancestral haplotypes were used for all subsequent analyses.

We summarize divergence among populations in several ways. Within a window, Ancestral Line Divergence (ALD) between two populations is the absolute difference in frequency, averaged over all ancestral lines:

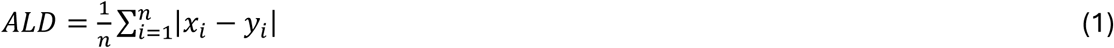

where *i* denotes ancestral line (1 to 187 in our study) and x and y the ancestral line proportions found in the two populations of the contrast (e.g. Ancestral versus evolved population 1). If the 187 frequencies are equivalent between two populations, ALD = 0. If we compare an idealized ancestral population (all ancestral lines with 1/187 proportion) to a descendant population where one ancestral line became fixed, ALD = (186(1/187) + 1(186/187))/187 ≈ 0.01. A second measure, “Ancestral Line Heterozygosity” (ALH) measures the amount of variation within a population. ALH is the probability that two randomly sampled alleles will be from different ancestral lines, *i.e.* the ‘expected heterozygosity’ at a locus with 187 alleles (Nei 1975). Finally, to evaluate the extent of hitch-hiking and test prediction of Fig 1, we calculated the correlation (r) of ancestral line frequencies between genomic windows at varying physical distances. This can be calculated separately for each ancestral line, but we use the average r across all 187 ancestral haplotypes as a summary statistic.

### Inference of selection

***—***Ideally, we would test for selection by comparing the likelihood of the data under competing models – a null hypothesis where the locus is neutral versus a more parameter rich model allowing selection to act in a treatment specific way. The resulting likelihood ratio statistic would then be compared to a chi-square distribution to determine significance. In this paper, we opt for a more conservative (and likely less powerful) non-parametric approach owing to our imperfect understanding of the “error distribution,” i.e. the statistical distribution describing how estimated haplotype frequencies differ from the truth. Preliminary analyses of our haplotype frequency estimates suggested a weak reference genome bias in mapping and a “flattening” of frequency estimates in genomic regions with lower read mapping fidelity. The latter happens when sequence reads from a single ancestor are split among multiple ancestors, those haplotypes that are similar in nucleotide sequence within a genomic window.

To address these concerns, we developed a test statistic from the multinomial probability distribution, but rely on permutation (and not asymptotic likelihood arguments) to assess significance. Within each genomic window, we calculate:

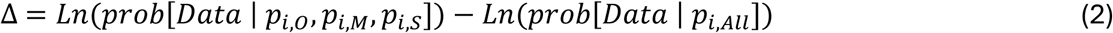

where *prob*[*Data* | *p̂*] is a multinomial probability taken over all populations and *p̂* is the vector of parameters:

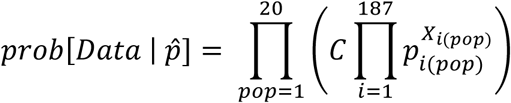

Here, C is the multinomial coefficient, *p*_*i*(*pop*)_ is the expected frequency of ancestral line *i* in population *pop* (i ranges from 1 to 187, pop from 1 to 20), and *X*_*i*(*pop*)_ is the observed count of ancestral line i for population pop within the window.

The terms in Δ are the log-likelihood under the alternative and null models, respectively. Each is calculated using (2) but with different specifications of *p*_*i*(*pop*)_from model parameters. For the null model, we specify that the expected frequency of each ancestral line is the same in all descendant populations. Thus, *p̂* is vector of 187 values (186 free parameters because they are constrained to sum to one). For the alternative model, we allow *p*_*i*_to vary by mating system – three values per line (*p*_*i*,*O*_, *p*_*i*,*M*_, *p*_*i*,*S*_) instead of one (*p*_*i*,*All*_). The optimal estimates are the simple averages of observed proportions across all populations (null model) or the averages across populations within each mating system treatment (alternative model).

Observed counts are over-dispersed relative to the multinomial even under the null model so we cannot apply the usual parametric approach (comparing 2Δ to the chi-square distribution with (3-1)*186=372 degrees of freedom). Instead, Δ provides a statistic that measures parallel change within treatments. Δ increases as the divergence among mating systems (in the full vector of *p*_*i*(*pOp*)_) increases relative to the variance among populations within treatments. We use whole genome permutation to assess significance. For each permutation replicate, we preserve the observed changes in ancestral line frequencies but permute by genomic location (window). We permute each population and then apply the testing regime to each window across the entire permuted dataset. We recorded the single highest value for Δ in each replicate and then repeated the process 5000 times. The permutation threshold is the 95^th^ percentile of this distribution. This is the standard approach of line-cross QTL mapping (Arends *et al*. 2010) and is substantially more conservative than FDR applied to individual tests (as is typical for GWAS).

Importantly, this is not a simple outlier approach because we are using the observed changes within populations to establish the null. The premise is that, with neutral evolution occurring independently within populations, there is no reason for coordinated changes to occur at the same locus across populations. Nor is there any reason why the direction and magnitude of these changes should be related to treatment. This analysis does not explicitly include the Ancestral samples. Instead, we use the Ancestral samples to evaluate the accuracy/bias of haplotype inference (see first section of **Results** below) and to ensure that ancestral lines started at approximately the same frequency. Equation (2) is easily generalized to include the ancestral population, but our interest here is the divergence among the mating system treatments. Finally, the raw output from HAFpipe is a continuous proportion and not a count for each ancestral line in each window. However, the underlying sample must be discrete: Each of the 400 alleles (200 diploid individuals sampled into each pool) is derived from a specific Ancestral sequence. For our selection tests that take counts as input, we obtained *X*_*i*(*pOp*)_for each line/population by multiplying the estimated proportion by 400 (the number of chromosomes in each pool) and then rounding to the nearest integer.

### Reanalysis of RNAseq data

***—***Brown and Kelly (2021) collected RNAseq data from flower buds on 151 of the 187 ancestral lines used in the current experiment. We remapped these sequence reads to the IM62v3 reference genome and quantified expression levels using Salmon v1.10.0 (Patro *et al*. 2017). After filtering (Program set 3 at https://github.com/jkkelly/Mating_system_evolve_resequence), we obtained expression levels for ca 25,000 genes across 277 plants (1-3 biological replicates for each of 151 of the ancestral lines). This is 3’ RNAseq data and we summarized expression level for each gene of each plant as Counts per Million (CPM) which was used as the response variable in downstream analyses. To assess the possibility that selected loci act through cis-regulatory changes, we tested all genes within each of the selected windows for association between expression level and response to selection. For each ancestral line *x*, we have a mean final frequency within the Outcrossed populations (*q̂*_*x*,*Out*_), a mean for Mixed-mating (*q̂*_*x*,*Mix*_), a mean for Selfing (*q̂*_*x*,*Self*_), and an estimated expression level (*Ê*_*x*_) for each expressed gene in the window.

Across all ancestral lines, we used the divergence of Mixed-mating from Outcrossing (*q̂*_*x*,*Mix*_−*q̂*_*x*,*Out*_) and Selfing from Outcrossing (*q̂*_*x*,*Self*_−*q̂*_*x*,*Out*_) as predictors of expression in a bivariate regression. If lines with relatively high expression increased in the Mixed-mating and Selfing populations, regression will yield positive slopes (and negative in reverse situation). For testing we used ranks instead of raw values for both dependent and independent variables because each variable routinely has outlier values (extreme lines for either expression response to selection). We applied a permutation to evaluate significance by randomly scrambling line mean expression levels against line identity (and thus response to selection). From each permutation replicate, we recorded the lowest p-value. The significance threshold is 1.0 x 10^-7^ because 95% of permutation replicates never produced a value this low. Gene annotation information (including homology to genes in Arabidopsis) was taken from Lovell *et al*. (2025) and gene expression in figures is reported as Ln(Counts per Million +0.5). To compare the genomic sequencing data from this study to that of Busch *et al*. (2022), remapped the pooled sequencing data from that study (four population to the IM62v3 reference genome. We then identified all SNPs segregating at intermediate frequency in the Ancestral lines, also called in Busch *et al*. (2022). We determine bi-allelic expected heterozygosity in all 26 samples at these SNPs and used these contrasts to assess final intra-population diversity in each population (Program set 4 at https://github.com/jkkelly/Mating_system_evolve_resequence).

## Results

We first compared the replicate ancestral population samples. Considering the overall frequency of each ancestral line across the entire genome, there is a very strong positive correlation between replicates (r = 0.93 between A1 and A2 in Fig 2A). The overall contribution of most lines is indistinguishable from the ideal value (1/187 = 0.0053) given that small deviations are inevitable due to varying DNA contributions to the pool. A1 and A2 were founded from different plants, DNA extractions, and library preps – the high correlation indicates nearly uniform per-line contributions. Notably, the overrepresented lines are the most sequence-similar to the reference genome (numbered points in Fig 2A). The reference genome used for mapping is based on ancestral line 62 and the other two lines (923 and Z537) have lower sequence divergence from line 62 than the other lines. Importantly, these lines are **not** key contributors to the major responses to selection (see below), but their elevated frequency (ca. 1.3% instead of 0.5%) suggests some bias in the inference of ancestral line frequencies. This bias is small on average, but it motivated our non-parametric testing procedure (see Methods), which admits imperfect inference of ancestral line frequencies so long as bias is unrelated to mating system treatment.

**Figure 2.**
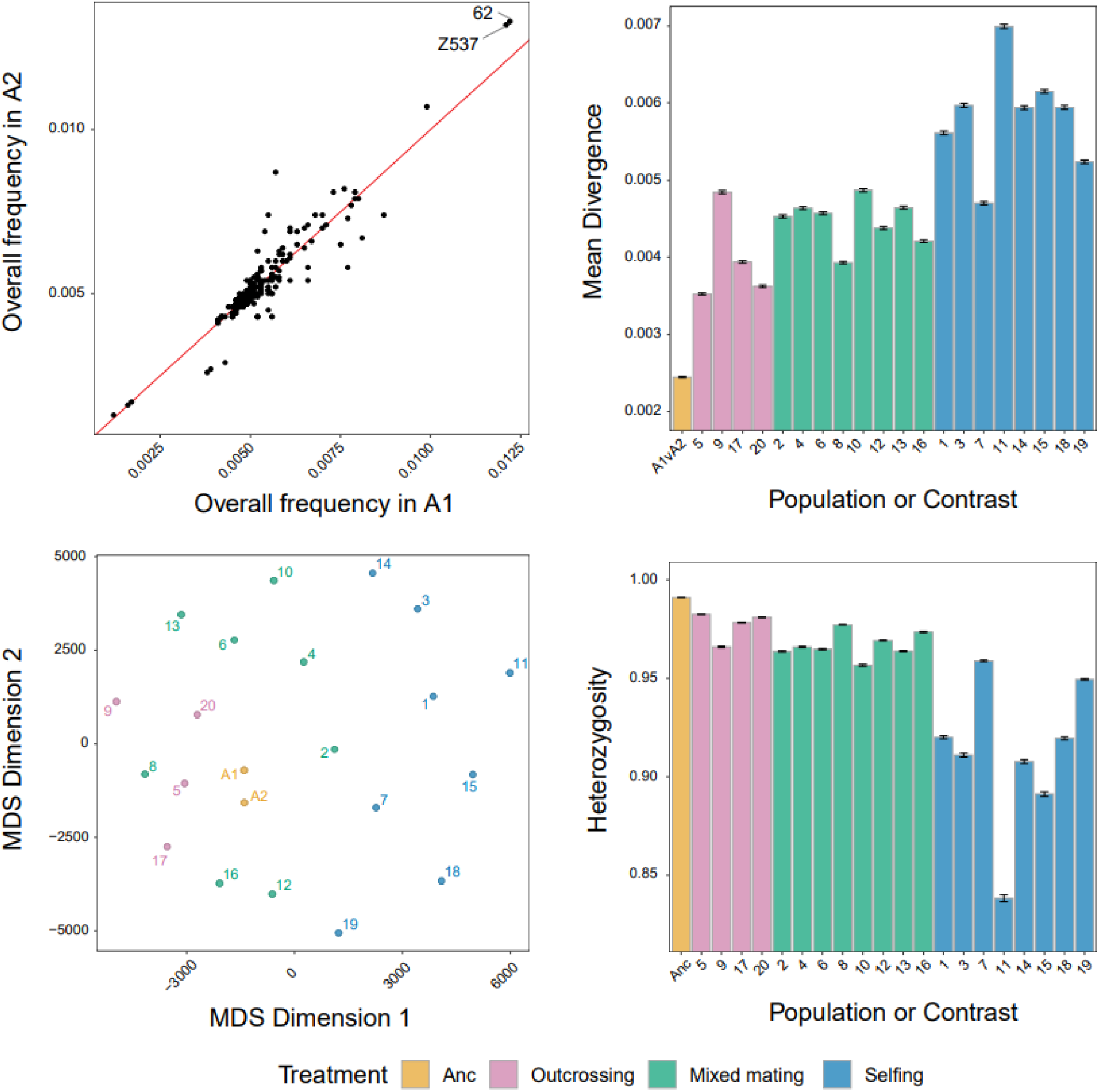
(A) The two ancestral pools show a very strong correlation in the overall frequencies of each ancestral line. (B) Divergence (mean ALS) differs among treatments. (C) A multi-dimensional scaling plot based on ALD illustrates differences among all 22 pooled samples. (D) Variation within populations (mean ALH) differs among treatments. In (B) and (D), “ANC” is the difference between A1 and A2, all other distances are between an experimental population (1-20) and the mean of A1 and A2.

Considering genomewide divergence of experimental populations from the Ancestral population (Fig 2B), ALD was lowest in the Outcrossing, intermediate in the Mixed-mating, and highest in the Selfing populations. There is significant variation among populations within treatments, particularly among Selfing populations. The overall pattern of divergence is illustrated by a multi-dimensional scaling plot based on the final ancestral line frequencies in each population (Fig 2C). Over most of the genome (although not the QTLs described below), the magnitude but not the direction of divergence is predicted by mating system. The Selfing populations became increasingly distinct from each other as well as from the Mixed and Outcrossing populations.

Inevitably, increased divergence of a population from the Ancestor is associated with reduced internal variation. The effect of mating system on expected heterozygosity (ALH) at the end of the experiment mimics the pattern for divergence (Fig 2D). ALH in the Ancestral population is near the theoretical maximum (>99%). The increase of some ancestral haplotypes and the loss of others over the ten generations reduces ALH, but only modestly in the Outcrossing populations. A pronounced decline occurred in several of the Selfing populations, particularly population 11. Population 11 was the only replicate where a single ancestral line approached fixation within a few regions of the genome.

The decline of ALH within each population from the ancestral value can be used to estimate the “effective population size”, N_e_, of each population. Noting ten generations of evolution (t = 10), we solved 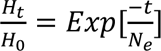 (Hartl and Clark 1989), for *N*_*e*_within each population (Supplemen*tal Table S1). Estimates* vary substantially by population, but the mean of Outcrossing populations (413) is almost twice mean of Mixed-mating (218) and 5.5 times greater than Selfing populations (74).

The striking reduction in variability within the inbreeding populations suggests an important effect of linked selection on the level of variation. Direct evidence for this is provided by correlated changes of ancestral line frequencies between genomic windows of varying physical separation (Fig 3). If a particular ancestral line carries a favorable allele, that ancestral haplotype should increase in the window that contains the relevant mutation. However, selection should also elevate neighboring windows which are likely to match the ancestry of the focal window owing to limited recombination since population founding. This correlation should decline with physical distance at a rate that depends on the mating-system specific rate of recombination. Since windows overlap (successive windows share 90% of their DNA), correlations are inevitable at small scales, even in the Ancestral population. However, Fig 3 shows that correlated changes are far greater in Selfing than Outcrossing populations at inter-window differences of 50kb to 1mb.

**Figure 3.**
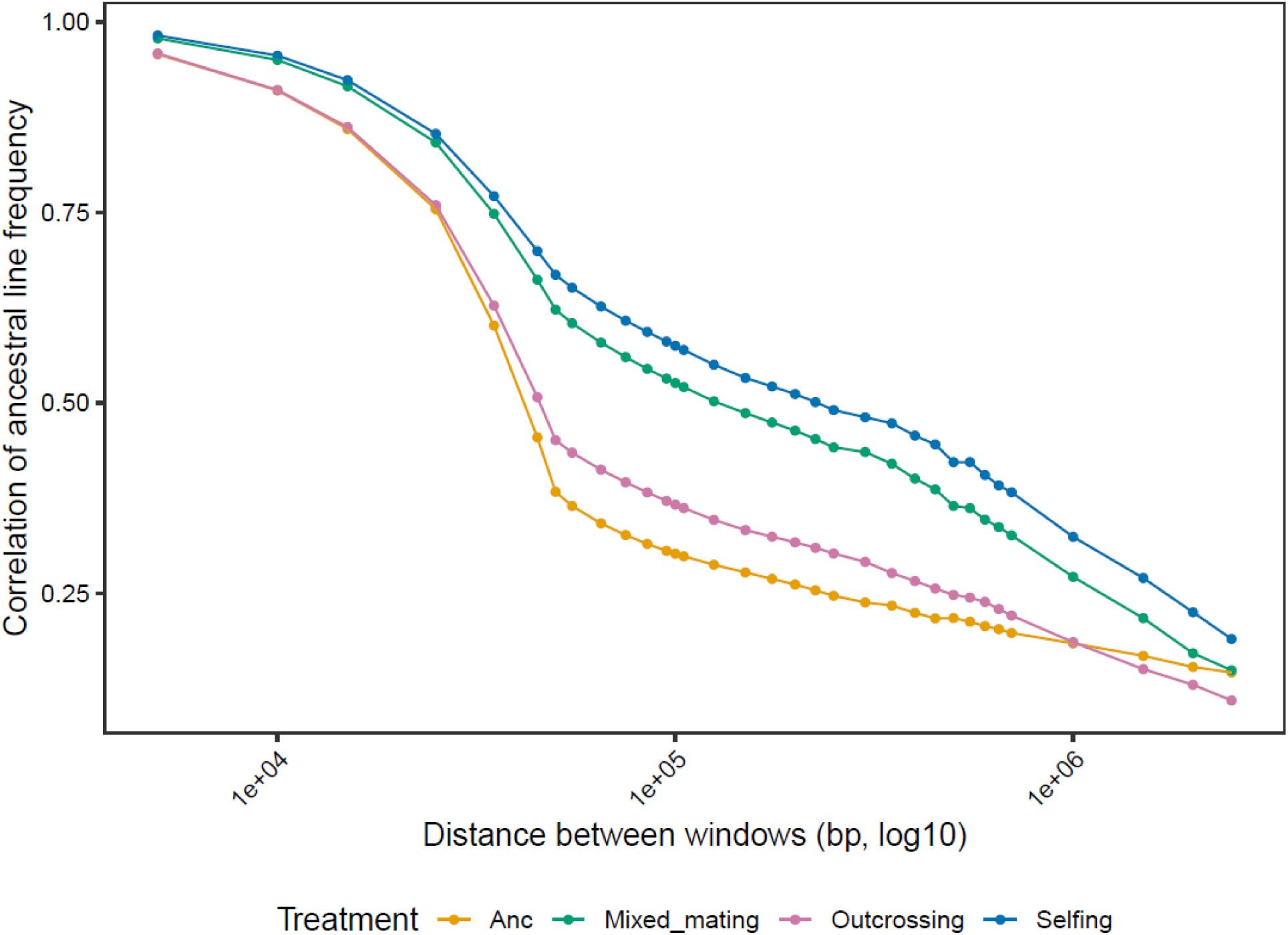
The correlation between ancestral line frequency estimates as a function of physical distance (log 10 scale) between windows is reported for each mating system treatment and the ancestral population.

Our test for mating system driven selection identified 1190 genomic windows where Δ exceeded the significance threshold (Fig 4A; Supplemental Table S2 reports tests for all 61886 windows). Many of these windows are overlapping and/or closely linked and reflect the same selective event. The strongest responses were on chromosome 6 (from location 0.5mb to 2.8mb), chromosome 12 (from 22.25mb to 25.5mb), and chromosome 1 (from 2.7mb to 6.75mb). The final ancestral line frequencies within the 1190 significant windows are reported in Supplemental Table S3. The specific ancestral haplotypes that increased within each window/treatment varied across QTLs. Ancestral line 750 had a high average frequency across significant windows in the Selfing treatment (9%), but this is mainly because it dominated the response within one very broad QTL on chromosome 1 (Fig 4A). The ancestral lines favored by mapping bias (62, Z537, and 923 in Fig 2A) are not elevated substantially within any treatment within QTLs.

**Figure 4.**
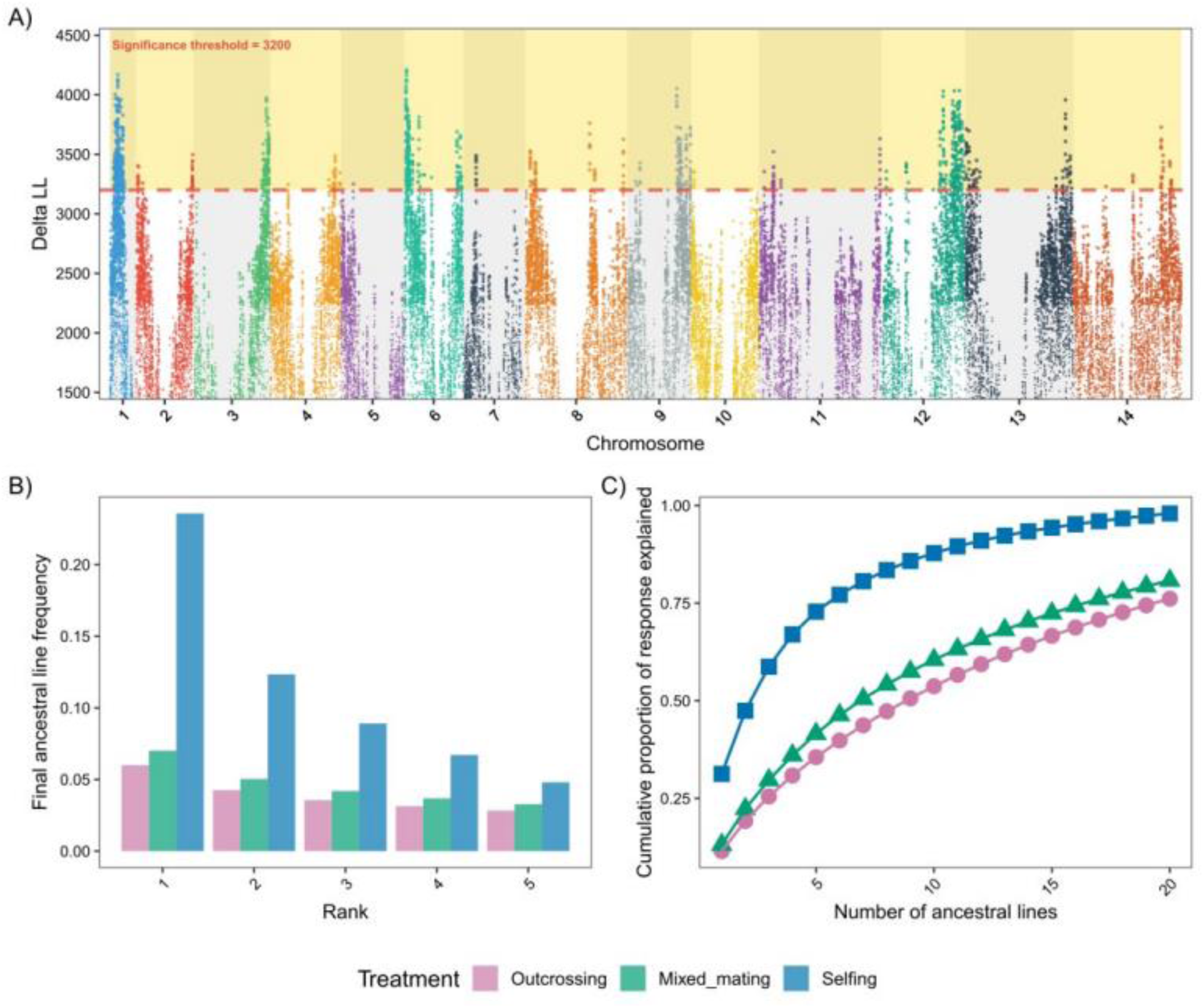
(A) Δ is reported for each window with the significance threshold as the horizontal broken red line. Chromosomes are distinguished by color. The gaps in the middle of each chromosome correspond to centromeric regions where Δ values were generally low. (B) The final frequency for the leading five ancestral lines (window specific) in each mating system treatment. (C) The proportion of the total response contributed by the first, second, third, etc., ancestral line within each treatment. Results in (B) and (C) are averages over all significant windows.

Most QTLs exhibit a strong response from initially uncommon alleles with within the Selfing populations – significance driven by 2-6 ancestral lines that increased substantially in this treatment. Oftentimes, Mixed-mating populations showed an elevated response in the same windows, in some cases via the same ancestral lines. However, there is much less overlap in terms of responsive ancestral lines between the Selfing and Mixed-mating than expected if the same selection regime was simply stronger in the Selfing treatment (explained more completely below). Of course, the pattern varies among selected regions. While less common, there were some windows where significance was driven by ancestral lines increasing substantially in the Outcrossing populations within minimal change in the other treatments, e.g. QTL at start of chromosome 13 (Fig 4A).

Within each significant window, we determined the average final frequency of each ancestral line in each mating system treatment (averaged across replicate populations) and then ranked them by magnitude of change from their initial starting frequency in the Ancestral (around 0.005). The “leading ancestral line” reached a final average of frequency of 24% in the Selfing treatment, but only 7% in the Mixed-mating treatment and 6% in the Outcrossing treatment (Fig 4B). If we sum the responses of the top five most responsive lines in each window/treatment, the corresponding proportions increase to 56% for Selfing, 23% for Mixed-mating, and 20% for Outcrossing. To some extent, these numbers reflect the fact that changes were generally greater in the Selfing populations, both within selected regions and across the genome (Fig 2B). However, the proportion of the total response that can be attributed to a few leading lines is much greater in Selfing than Outcrossing or Mixed populations (Fig 4C). In the Selfing treatment, the first five ancestral lines explain about 73% of the total response, on average. We need to include the top 16 to reach that proportion in the Mixed-mating treatment and 19 for the Outcrossing treatment. Supplemental Table S4 reports the top ten responsive ancestral lines within each significant window within each mating system treatment.

Within most QTL, the ancestral lines increasing in the Mixed-mating populations were different from those that increased in the Selfing populations. Fig 5 classifies the responsive lines (the top 10 within each significant window within each mating system) according to whether they are unique to treatment or shared across treatments. In most windows, overlap between treatments was limited. For every ancestral line that increased in both the Selfing and Mixed-mating treatments, there were about four that only increased in the Selfing (27.8/6.8=4.1) and nearly the same number that increased only in the Mixed-mating treatment (25.7/6.8=3.8). The Outcrossing populations also show low overlap in terms of leading lines with the other treatments, but this is expected given that the Ancestral mating system is predominantly outcrossing. Many of the top 10 responsive lines in the Outcrossing treatment had final average frequencies of only 1-2%. Increases of this magnitude could be driven at least partly by drift. Lack of overlap is expected with drift.

**Figure 5.**
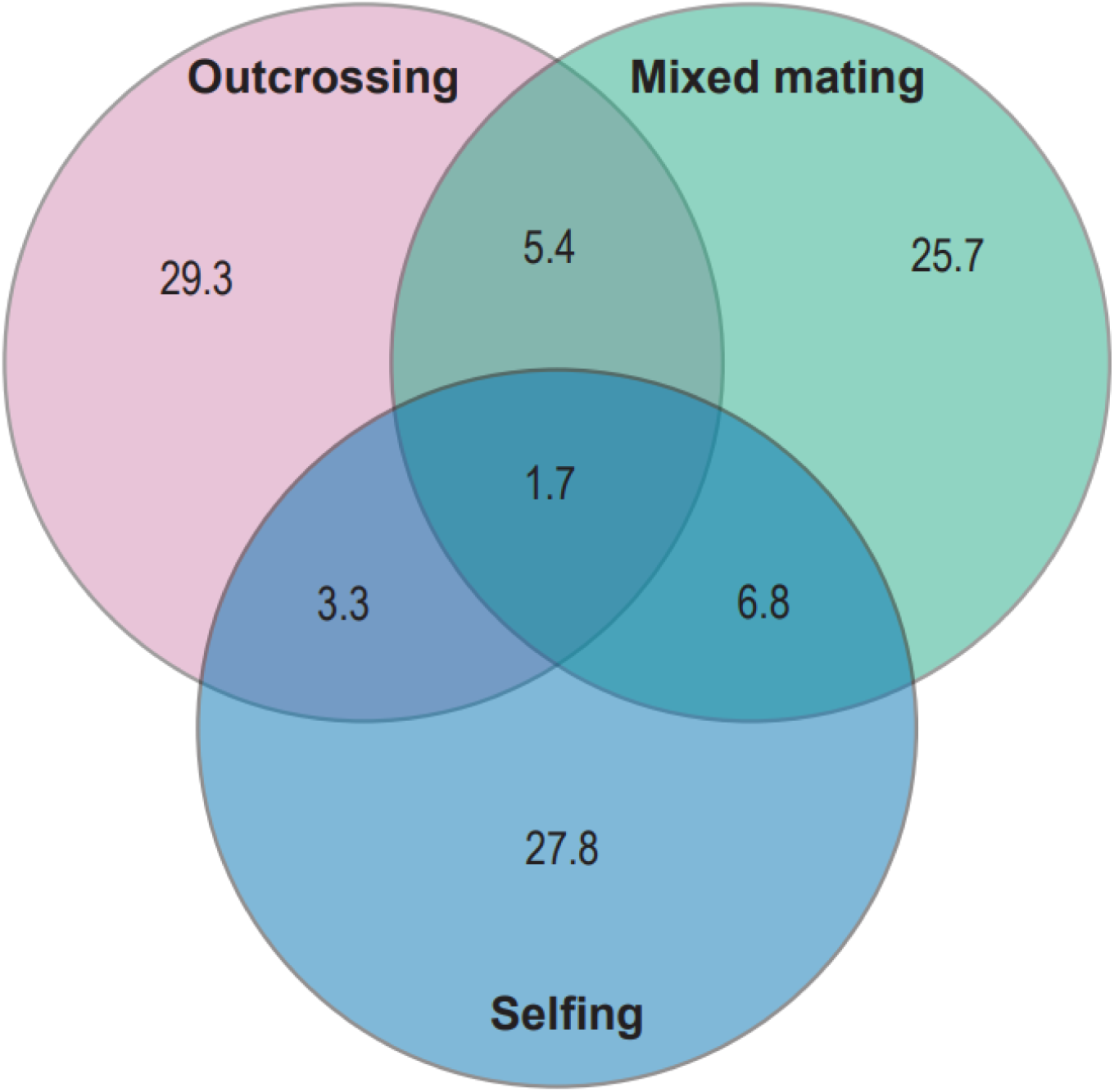
The overlap of the top 10 responsive lines by mating system treatment across all significant windows. The numbers are percentage of ancestral lines in each category.

We next tested whether response to selection was associated with differential gene expression within significant windows (QTLs), evaluating the idea that cis-eQTL are responsible for fitness effects. We found 79 genes where expression was significantly associated with the evolutionary responses of the ancestral lines (Table 1). Remarkably, there is a very strong tendency for ancestral lines that increased in the Selfing populations (relative to the Outcrossing populations) to exhibit reduced expression at significant genes: 57 of 79 genes are downregulated (Sign test p < 0.001; Spearman correlations in Table 1). In contrast, these same genes show no directionality in the ancestral lines that increased in the Mixed-mating treatment (37 downregulated, 42 upregulated).

The tests in Table 1 are based on gene expression in flower buds from 151 of the 187 Ancestral lines. Because expression estimates are specific to ancestral line, they can be compared to the evolutionary changes in those same lines across populations in each mating system treatment. There was no need to bin the lines *a priori* into functionally distinct “expression alleles” for testing – locus specific responses determined significance. However, we can bin ancestral lines *a posteriori* given the observed pattern of changes within a genomic window in order to characterize the varied relationships between gene expression and response to mating system treatment across the genome. Figures 6-7 illustrate four of the differentially expressed genes in Table 1, two from within each of the two most significant selection peaks identified by Δ (on Chromosomes 1 and 12, respectively). While the direction of differential expression (up in the Selfing treatment) may be opposite the overall trend, these cases correspond to strong candidate genes (see Discussion) and illustrate how closely linked genes can have importantly different patterns.

**Figure 6.**
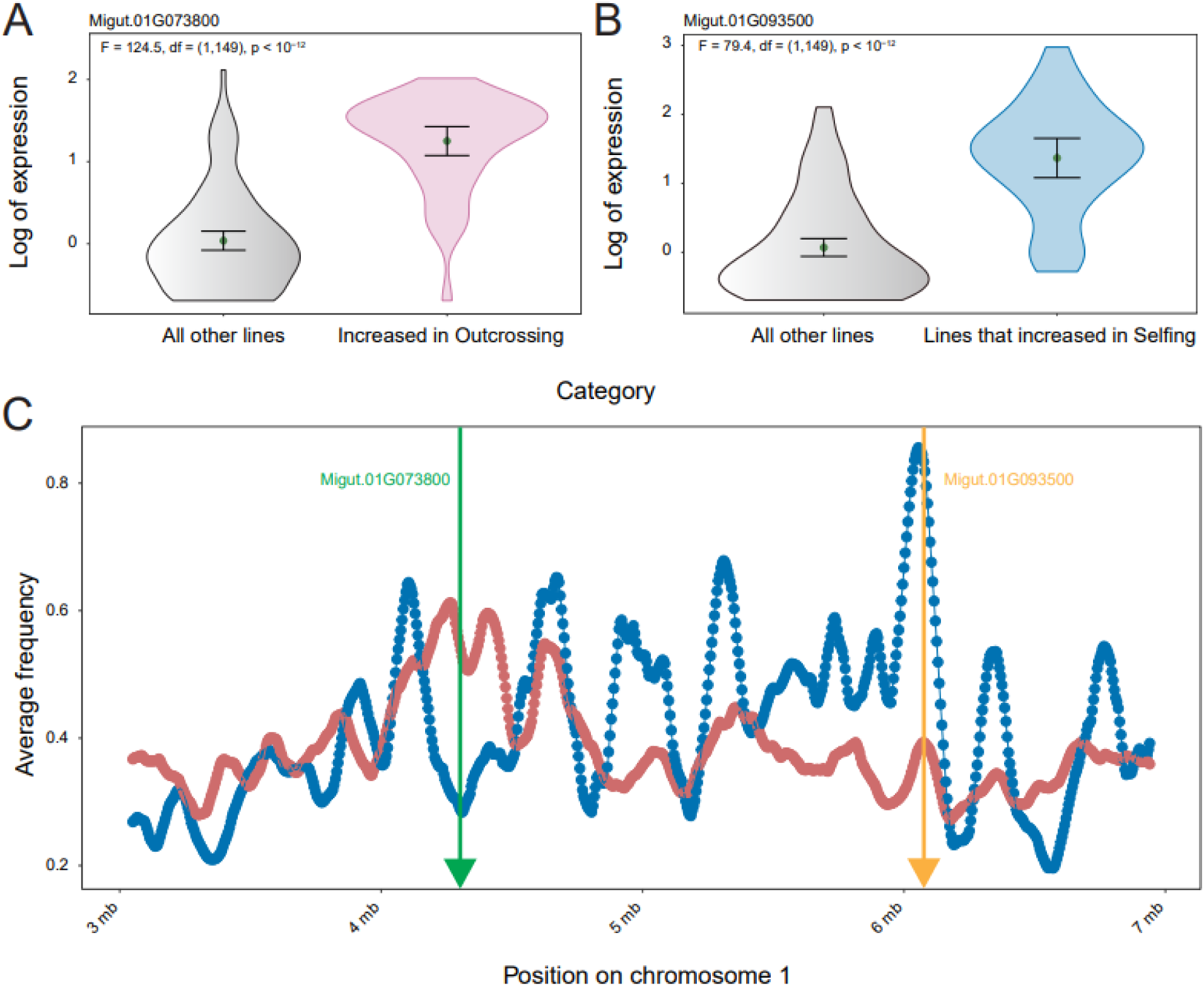
Two of the genes on chromosome 1 most significant for differential expression are (A) Migut.01G073800 and (B) Migut.01G093500. (C) The trajectories for the responsive line sets within each QTL: Blue = 37 ancestral lines that increased in the Selfing populations within the significant window for Migut.01G093500, Red = 49 ancestral lines that increased in the Outcrossing populations within the significant window for Migut.01G073800. Responsive lines are those where the final average frequency across populations with treatments was greater than 0.01 in the relevant treatment/window.

**Figure 7.**
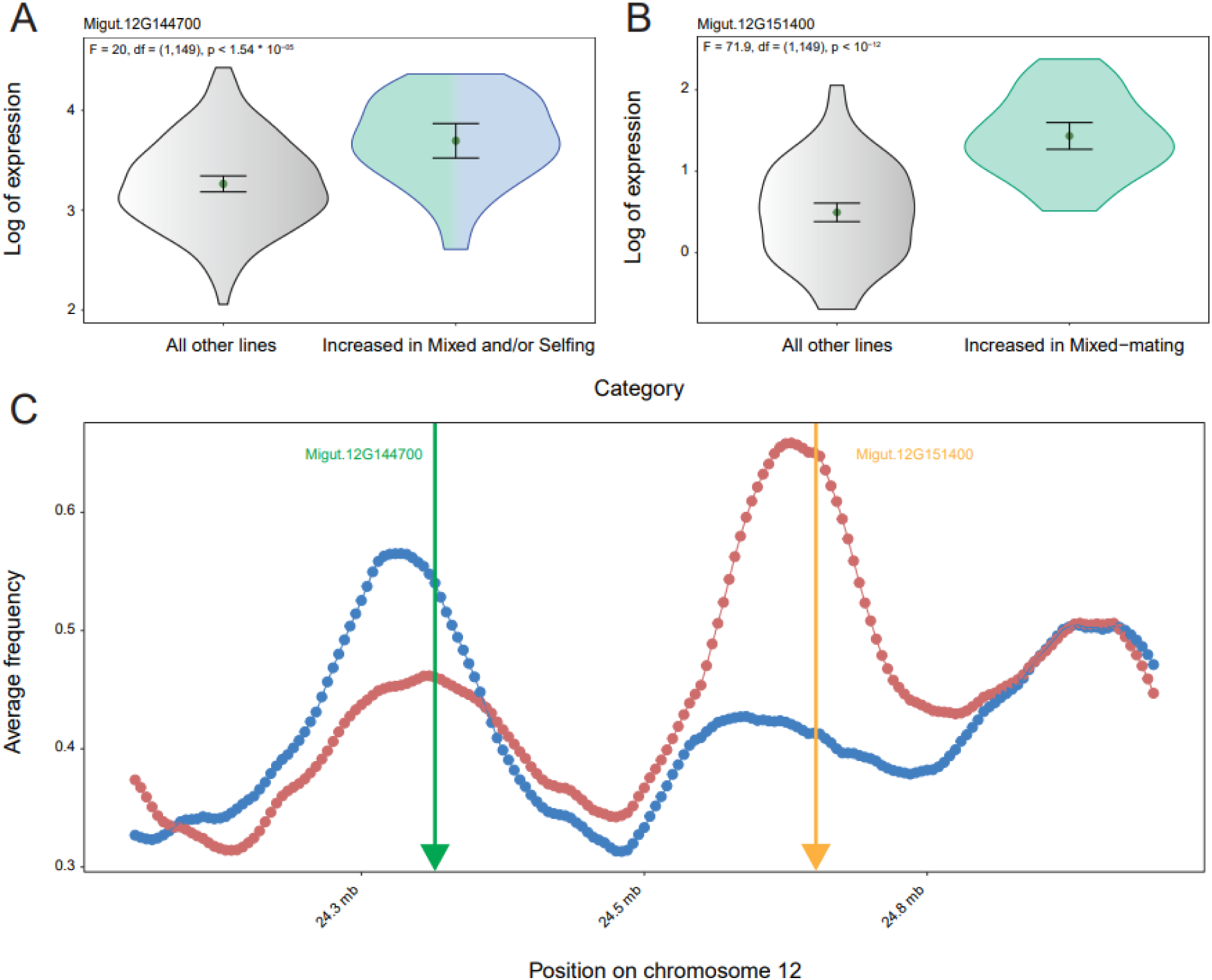
Two of the genes on chromosome 12 that are most significant for differential expression are (A) Migut.12G144700 and (B) Migut.12G151400. Other conventions the same as in Fig 6.

The broad QTL on chromosome 1 is generated mainly by six ancestral lines that showed elevated average frequencies in the Selfing populations in most of the windows over a 4mb span (Fig 4A, left). Another 30 or so of the ancestral lines show more localized increases within the Selfing treatment populations over this portion of the genome. More specifically, in the window centered on 6.1mb, 37 of the ancestral lines obtained average final frequencies of greater than 1% across the 8 selfing populations (cumulatively about 85% as indicated by the blue trajectory in Fig 6C). 32 of these 37 lines were measured for gene expression. They express the gene Migut.01G093400 (Fig 6B), a lectin protein kinase, at an average level nearly 4-fold higher than the other 119 ancestral lines measured for expression. Like the original test based on rank correlations (between expression and response, p = 10^-11^ in Table 1), a simple ANOVA based on this dichotomous classification of lines is highly significant (p < 10^-12^, Fig 6B).

The trajectories of Fig 6C track the frequencies of ancestral lines that increased in one or more treatment owing to selection – this is hitch-hiking directly observed. The trajectories fluctuate owing to the history of recombination events within each replicate. Importantly, the cumulative change of the “blue” ancestral lines within the Selfing populations varies over the broad QTL on chromosome 1. A window upstream (at 4.3mb) shows a qualitatively different pattern where a large group of ancestral lines increased in the Outcrossing populations to a cumulative frequency of 62% (red trajectory in Fig 6C). A gene in this window (Migut.01G073800 in Fig 6A) is upregulated among the Outcrossing responsive lines, about 3.3x higher than in the remaining non-responsive lines. The pattern of gene expression differentiation—Outcrossing divergent in Fig 6A versus Selfing divergent in Fig 6B—occurs because the underlying patterns of ancestral line frequencies changes between these two loci despite generally high values for Δ (Fig 4A).

Figure 7 illustrates two genes on chromosome 12: Migut.12G151400 (Fig 7A) is an Ankyrin repeat family protein located about 350kb upstream of Migut.12G144700 (Fig 7B), a nucleic acid-binding OB-fold-like protein. Both genes show differential expression, but with different underpinnings. In genomic window containing the Ankyrin repeat gene, a strong response is observed with ancestral lines increasing in both the Mixed and Selfing treatments. Those ancestral lines that increased in these treatments are upregulated relative to non-responders (Fig 7A). The pattern of ancestral line changes substantially between this gene and the closely linked Migut.12G144700 (Fig 7B). The ancestral lines that were elevated in the Mixed-mating populations in the Migut.12G151400 window were also elevated in the Migut.12G144700 window. However, a larger collection of ancestral lines joined the Mixed-mating response in the latter window. This produces a much stronger signal of selection driven by the Mixed-mating treatment (see red trajectory in Fig 7C). Between these two windows, the Selfing response became decoupled from the Mixed-mating response. At Migut.12G151400, the lines that increased in Mixed-mating (relative to the Outcrossing) populations had elevated expression (Sperman r = 0.32 in Table 1) while the lines that increased in the Selfing populations had lower expression (Spearman r = −0.41). In this small genomic region, the evolutionary responses of the Mixed-mating and Selfing treatments were decoupled.

## Discussion

### The genes under selection

***—***Mating system loci are valuable targets for genetic mapping. In flowering plants, genes that alter the mating system, say by changing flower color or by allowing individuals to self-fertilize, can immediately generate pre-zygotic isolation among populations or lineages. The phenotypic effects of these loci are often recognized as species defining characteristics (Ortiz-Barrientos 2013). Mating systems illustrate how genetic complexities such as allelic dominance and effect size can qualitatively alter evolutionary outcomes. Rare, recessive variants are largely invisible in outbreeding populations but generate substantial fitness variation in populations that reproduce partially or fully by self-fertilization. This is true both when (partially) recessive alleles are advantageous (Caballero and Hill 1992; Charlesworth 1992; Bachmann *et al*. 2019) and when they are deleterious (Lande and Schemske 1985). Mutations that cause a major increase in selfing rate can sometimes increase when high inbreeding depression prevents evolution by small steps (Uyenoyama *et al*. 1993; Kelly 2005). Mapping results are most compelling when genetic details can alter the outcome of phenotypic evolution.

Figures 3–7 illustrate both the potential and the challenges of using experimental evolution to map mating system modifiers. Many of the genes in Table 1 are excellent candidates as effectors of selfing rate. For example, Migut.01G073800 (Fig 6A) is homologous to the CCR4-associated factor 1a (CAF1a) of *Arabidopsis thaliana*. CAF1a is a component of the CCR4–NOT complex that regulates mRNA (Behm-Ansmant *et al*. 2006). Experimental overexpression of CAF1a alters the rate of maturation to flowering and the expression of several canonical flower time genes including *FLOWERING LOCUS T (FT)*, *SUPPRESSOR OF OVEREXPRESSION OF CO 1* (*SOC1*), *EARLY FLOWERING3 (ELF3)* and *FLOWERING LOCUS C* (*FLC*) (Lv *et al*. 2023). In the present study, differential expression of Migut.01G073800 was driven by an evolutionary response in the Outcrossing populations (Fig 6C,left). One of the strongest phenotypic responses in the Outcrossing treatment was accelerated flowering (see Fig 3 of Tusuubira and Kelly (2024)).

Migut.02G051900 on Chromosome 2 showed divergent responses in terms of gene expression between Mixed-mating and Selfing treatments (Table 1). This gene produces a Mitogen-Activated Kinase Kinase Kinase (MAP3K) protein. MAP3K genes are critical components of the intracellular signaling cascades that regulate many developmental processes in plants (Chen *et al*. 2015; Xie *et al*. 2023). In Arabidopsis, MAP3K genes affect flower development (Cho *et al*. 2008) and alter the relative sizes of floral organs through differential cell proliferation (Meng *et al*. 2012). Jia *et al*. (2016) demonstrated effects of the entire MKK7-MPK6 cascade, which includes a MAP3K protein, on both shoot branching and filament elongation. Both phenotypes differ among mating system treatments with filament length (the stalk that supports the anther sacs) a critical determinant of stigma-anther separation. Stigma-anther separation is a strong morphological predictor of self-seed set in *M. guttatus* (Fishman and Willis 2008; Bodbyl Roels and Kelly 2011).

Perhaps the most interesting candidate is Migut.01G093400 (Fig 6B) which is homologous to the S-locus lectin protein kinase of Arabidopsis (AT4G27290). This gene is one of two components of the SRC/SRK Self-Incompatibility system in the plant family Brassicaceae (Nasrallah and Nasrallah 2014) and lectin receptor kinases have been shown to control self-pollen recognition in other families, e.g. Burgin *et al*. (2025). *M. guttatus* does not have a genetic self-incompatibility system, but lectin protein kinases are generally involved in cell-to-cell signaling and pollen-pistil interactions (Bellande *et al*. 2017). Quantitative changes to stigma receptivity over the course of the floral lifespan are likely essential to effective self-pollination in *M. guttatus* (Arathi *et al*. 2002). Other lectin protein kinases have been shown to alter pistil length (Xiao *et al*. 2021) even pollen viability (Wan *et al*. 2008). The *M. guttatus* reference genome has three duplicate copies of AT4G27290 located consecutively over 35kb of chromosome 1 (Migut.01G093400, Migut.01G093500, and Migut.01G093600). Two of the three copies show strong up-regulation in the ancestral lines that increased in the Selfing treatment (Table 1).

Figs 6-7 also illustrate how a fundamental feature of selection with inbreeding hinders fine scale genetic mapping. Increased homozygosity reduces the efficacy of recombination and so hitch-hiking will occur over larger genomic regions (Fig 3) and QTL peaks driven by responses in the Selfing populations are thus quite broad (Fig 4A). The gene expression data provides much finer resolution (physically to the gene scale) and can be used to identify the portions of broad QTLs that are most likely to contain the actual targets of selection (Table 1). Unfortunately, most broad QTLs contain several genes that are highly significant for differential expression. Many of genes in Table 1 have *Arabidopsis* homologs that have been subjected to functional manipulation with consequent measurement of altered expression effects plant traits. The complication is that while our candidate genes are implicated in response to selection, they are not linked to specific phenotypes and many traits changed in response to mating system treatment (Tusuubira and Kelly 2024). Gene expression measured in flower buds is plausibly related to the phenotypic changes that occur with mating system shifts (Slotte *et al*. 2013), but follow-up studies are required to link specific genetic loci to specific phenotypic effects.

The strongest inference from Table 1 is not based on any particular gene. Across all 79 differentially expressed genes, almost three quarters were downregulated in ancestral lines that increased in the Selfing treatment populations. Comparing several selfing species of *Capsella* to an outcrossing sister species, Zhang *et al*. (2022) found that differential regulation was highly asymmetric: Downregulation was twice as frequent as upregulation in the selfing species. A gene expression asymmetry between divergent species could be explained by evolution within lineages after they transitioned to predominant self-fertilization. If mutations in cis-regulatory regions are more likely to reduce than increase expression, selfing species may show an abundance of down-regulated genes because mildly deleterious alleles are more likely to fix with selfing (Charlesworth and Wright 2001). An alternative explanation is that down-regulation is simply more frequently required to produce the phenotypic changes necessary for successful reproduction by selfing. In our study, the asymmetry must be due to the recruitment of standing variation and parallel increase of alleles in multiple replicate populations indicate adaptation and not the fixation of *de novo* deleterious mutations. Excepting the caveat described below, these results favor the adaptive downregulation hypothesis.

### The dynamics of allele frequency change with inbreeding

***—***Theoretical studies have produced an array of predictions regarding the micro-evolutionary consequences of inbreeding. Inbreeding fundamentally changes the way that genetic variation translates into phenotypic variation and thus how natural selection “sees” the alleles that segregate in populations. At individual loci, increased homozygosity can facilitate adaptive change when favorable alleles are recessive (GlÉmin and Ronfort 2013) and reduces inbreeding load by ‘purging’ deleterious alleles (Charlesworth and Charlesworth 1987). For quantitative traits, inbreeding alters the resemblance among relatives (Cockerham 1983; Cockerham and Weir 1984) and how trait means change as a function of genetic and fitness parameters (Kelly 1999a; Kelly 1999b). Inter-locus associations, both identity disequilibrium and linkage disequilibrium (LD), take on greater importance with inbreeding. The recombination inhibiting effect of selfing can be detrimental if favorable alleles become trapped in different competing lineages within a population (Hayashi and Ukai 1994), although this same process can also preserve variation within selfing populations (CLO *et al*. 2020). Selfing can help to preserve favorable gene combinations when loci interact to determine fitness (Allard 1975). Over longer time scales, mating system can substantially alter genome-wide patterns of polymorphism and divergence (Glemin 2007; Hartfield *et al*. 2017).

Using fully sequenced lines as founders provides an empirical lens on micro-evolution under different mating systems (Fig 1). One of the hypothesized effects of selfing – that selection should generate pervasive hitch-hiking – is immediately evident from the observed frequencies of ancestral haplotypes after only ten generations of evolution (Fig 3). There is much stronger covariation in allele frequency change between loci separated by 10kb to 1mb in the Selfing and Mixed-mating populations than in the Outcrossing populations. This is a consequence of both selection and variably effective recombination. Among the ancestral lines, LD among SNPs declines to near zero at intergenic physical distances (20-50kb, Supplemental Table 2 in Kelly (2022)). This means that the hitch-hiking evident at greater distances in Fig 3 is driven by LD that emerges due to selection as favored ancestral lines carry long haplotypes to high frequency. Given the recombination map of *M. guttatus* (Flagel *et al*. 2019), chromosomes in founding individuals should be a mosaic of two ancestral lines on average. Given the physical size of *M. guttatus* chromosomes, ancestral haplotypes should have averaged about 10mb in size at the onset of selection (Generation 0 in Fig 1). Recombination fractured these haplotypes over the course of the experiment, although much more finely in the outcrossing populations. The patterns evident in Figs 6-7 suggest the interesting possibility that selection may have favored multi-locus haplotypes within different mating system treatments, each constituted from different allelic combinations at linked loci.

Selfing populations exhibited the most substantial evolutionary change, both within QTL (Fig 4B) and genomewide (Fig 2). Amplified stochastic change is predicted with inbreeding owing to both genetic drift and linked selection (Robertson 1961; Pollak 1987; Caballero and Santiago 1995). Busch *et al*. (2022) recently analyzed pooled sequencing from the prior study of Bodbyl Roels and Kelly (2011) where replicate populations of *M. guttatus* were maintained for ten generations with or without bumblebee pollinators. They demonstrated dramatic declines in SNP diversity within the “no-bee” populations relative to “bee” populations. We reanalyze that data alongside results from the present study, scoring SNP diversity in a comparable way (same reference genome, filters, etc.). Despite differences in experimental design and adult population sizes, final SNP diversity in the no-bee populations of Busch *et al*. (2022) is very similar to the Selfing treatment populations of this study. Both studies indicate reductions in variability with selfing that greatly exceed a 50% reduction in effective population size expected from drift without linked selection (Supplemental Table S1).

Figure 4 provides a view of the “allele frequency spectrum” for mating system modifiers that segregate in outcrossing populations. Allele frequencies at SNPs across the genome are very strongly correlated between the sequenced ancestral lines and direct collections from the wild population (r = 0.97 reported in Kelly 2022). In QTLs driven by evolution within the Selfing treatment, 5-10 ancestral lines typically generated most of the response (Fig 4C). If these lines all carried the same advantageous allele, initial frequencies for selfing-favorable alleles would typically be 3-5%. In contrast, increases in the Mixed-mating and Outcrossing treatments were distributed over a much greater number of ancestral lines (each of which increased to a lesser extent over the course of the experiment). The initial rarity of alleles favorable to selfing may contribute to the genomewide diversity pattern. The rapid increase of rare alleles has a diversity-reducing effect similar to that of a hard sweep (Kaplan *et al*. 1989; Hermisson and Pennings 2005), except here extending over a much larger genomic regions owing to reduced recombination. In a study of contemporary evolution in a highly selfing barley species, Landis *et al*. (2024) observed near fixation of large haplotypes that were rare in the founding population. The novel experimental environment caused a rapid increase of a specific barley haplotypes, with consequent effects on genomewide variation.

How frequently do outcrossing population harbor low-frequency genetic variants that are favorable if pollinator service declines and selfing is required? Raduski *et al*. (2012) surveyed data from genetically Self-incompatible plants; species that are often considered ‘obligately outcrossing.’ They found that if enough individuals are scored within SI populations, one or a few individuals are routinely found to be partially self-compatible. If this self-compatibility has a genetic basis, the underlying alleles may have similar population frequencies as the selfing-favorable alleles mapped here.

A potential difficulty with these arguments: We are equating the number of ancestral lines that increased with the number that are carrying a self-favorable allele. It is possible that all carriers are *not* competitively equivalent. Inbreeding depression is severe within the IM population of *M. guttatus* (Brown and Kelly 2020). All the ancestral lines carry deleterious alleles, most of which are at least partially recessive (Willis 1999b; Willis 1999a). Lines that happen to have deleterious recessives linked to the positively selected locus will be a disadvantage relative to lines that do not, even if the latter lines have equal (or more) deleterious load across the genome. A shift to complete selfing transforms a population into a collection of competing lineages. Through a combination of chance or selection, one (or a few) lineages might capture a specific genetic background, one with fewer deleterious alleles and/or a collection of favorable alleles, and then outcompete other lineages.

We included the Mixed-mating treatment in this study in part because the inclusion of a low rate of outcrossing can substantially alter the evolutionary dynamic associated with compulsive selfing. Occasional outcrossing can effectively combine advantageous alleles that otherwise would be isolated in competing lineages into the same genotype. Even with only 10% outcrossing, our Mixed-mating populations exhibited whole genome patterns more similar to the Outcrossing populations than to the fully Selfing populations (Figs 2,4B,4C). Theoretical studies of many population genetic processes have shown that a little bit of outcrossing can go a long way (Maynard Smith 1978). Again however, we have to acknowledge the possible contribution of inbreeding depression. While only 10% of seed came from outcrossing, outbred plants might have constituted a much greater proportion of the adult population in the Mixed-mating populations owing to superior vigor (Scofield and Schultz 2006).

Mixed-mating is a very common mating system in flowering plants (Vogler and Kalisz 2001). In nature, outcrossing populations will often experience a partial but not complete loss of pollinators (Bishop *et al*. 2023; Acoca-Pidolle *et al*. 2024) and the Mixed-mating treatment provides a better treatment of this scenario than the complete Selfing treatment. We did not anticipate qualitatively distinct outcomes between Mixed-mating and Selfing in terms of the identity of the ancestral lines responding to selection. The lack of overlap evident in Fig 5 suggests the interesting possibility that partial and complete pollinator loss may actually favor different alleles at the same multi-allelic loci.

### Conclusion

***—***We developed a novel testing framework using ancestral haplotype inference to identify mating system loci in *Mimulus guttatus*. As expected from theory, we found that Selfing populations have increased homozygosity, pervasive hitch-hiking, and elevated stochastic change in allele frequencies relative to partially or fully outcrossing populations. Despite the latter, about 20 regions of the genome show parallel evolution within mating system treatments. These QTLs are usually quite broad (often greater than 1mb) but we could resolve candidate genes within QTLs using RNAseq data from the ancestral lines. In several cases, we found closely linked candidate genes, each of which may have been causal to the selection response. These results support the hypothesis that inbreeding may allow selection on gene combinations through its inhibitory effects on recombination. Individual based studies will be required to establish causal relationships between genotype, gene-expression, and reproductive strategy. Finally, we found a general tendency for down-regulation of candidate genes in the Selfing populations, a result that mirrors transcriptome differences that have been documented between established outcrossing and selfing sister species. Gene expression appears to be an important component of the “selfing syndrome.”

## Data availability

The pooled sequencing data from experimental populations are being submitted to NBCI. Other sequencing data can be downloaded directly from NCBI though accession information in the cited papers. All python programs used for analysis are available at https://github.com/jkkelly/Mating_system_evolve_resequence.

## Supplemental Materials

**Table S1. The estimates for effective population size, Ne, of each population obtained from Ancestral Line Heterozygosity (ALH).**

**Table S2. The testing results are reported for all windows. LL0 and LL1 are the log-likelihood scores for the null model and alternative model, respectively. Δ is LL1-LL0. “Sig” indicates whether the test exceeds the genomewide significance threshold of 3200.**

**Table S3. The average frequency of each ancestral line in each treatment across the 1190 significant windows.**

**Table S4. The top ten most responsive ancestral lines (highest average final frequency) are reported for each significant window for each treatment.**

